# Assessment of Tuberculosis incidence and treatment success rates of the indigenous Maká community in Paraguay

**DOI:** 10.1101/620161

**Authors:** J. Fröberg, V.G. Sequera, A. Tostmann, S. Aguirre, C. Magis-Escurra

**Affiliations:** Radboud University Medical Center, Center of infectious diseases-TB expert center Dekkerswald, department of respiratory diseases, Nijmegen-Groesbeek, the Netherlands; Health Surveillance Department, Ministry of Health, Asunción, Paraguay; Department of Primary and Community Care, Radboud Centre for Infectious Diseases, Radboud university medical centre, Nijmegen, The Netherlands; National Tuberculosis Control Program, Asunción, Paraguay

**Keywords:** Epidemiology, aborigines, outcome, childhood tuberculosis, surveillance

## Abstract

**Setting:** In Paraguay, 1.8% of the population are indigenous people. The Maká community mainly live in urbanized areas in the Central Region. This study focuses on the epidemiology of tuberculosis (TB) among indigenous Maká and the non-indigenous people living in the Central Region, the biggest metropolitan area of the Paraguay.

**Objectives:** This study aims to analyze the TB incidence and treatment success rate of the urbanized Maká indigenous population

**Design:** Retrospective cohort study of 6,147 registered TB patients with 387 Maká indigenous people, from 2005-2017.

**Results:** Compared to the non-indigenous population in the Central Region, the Maká had a 66 times higher TB incidence, a lower median age at diagnosis (3 vs. 33 years; P<0.001), less bacteriological diagnosis (55.0% vs. 77.8%; P<0.001), and a higher treatment success rate of 75.2% vs. 67.8%. Directly observed therapy coverage was higher among the Maká (89.4% vs. 47.1%; P<0.001).

**Conclusions:** The Maká showed a disproportionately high TB incidence in children. Treatment success rates did not reach the WHO standards of 85%. If the diagnosis in children from this period can be confirmed, the public health system should intensify their focus on the Maká, increasing case finding and contact tracing activities in the whole population.

## INTRODUCTION

In Paraguay, a country with a population of 6.8 million, each year approximately 2,800 people are diagnosed with active tuberculosis (TB).(1) In 2016, TB mortality was estimated at 270 people (9%).(1) The indigenous populations of Paraguay, forming only a small part of the total population (1.8%)(2) are more vulnerable for TB. The national TB-burden in Paraguay is classified as intermediate with 42 TB cases/100,000 inhabitants, whereas the burden for indigenous people was reported at 272/100,000 in a National survey in 2014.(1-5) Since 1985, the indigenous Maká population (Maká), one of 19 tribes in the country, mostly lives in urbanized areas (77.4%), mainly Mariano Roque Alonso (MR Alonso) and Villa Hayes(2), which are part of the Central Region; the biggest metropolitan area in Paraguay. Successful treatment is essential for TB-control. If the recommended TB treatment regimen(6) is not completed, the chance of recurrence, TB transmission, and development of acquired drug resistance is high.(7-9) To improve treatment adherence in Paraguay, Directly Observed Treatment Strategy (DOTS) was introduced in 2002. Research on the DOTS coverage and the improvement of treatment outcomes in Paraguay is still lacking. Furthermore, the differences between the indigenous and non-indigenous people have never been thoroughly analyzed.

The current study focused on the Maká living in the Central Region of Paraguay.(2) By evaluating the patient demographics of the Maká and the non-indigenous living in the Central Region, assessing current TB treatment success rates, and analyzing the registration performance of the National TB program (PNCT) in this region, we hope to provide useful information that surpasses the surveys currently performed in Paraguay. This knowledge will facilitate more focused interventions for TB-control in the Central Region and improve registration by the PNCT to track the progress of these interventions.

## METHODS

### Study design and study population

In this retrospective cohort study from 2005-2017, TB registration data was analyzed from the Central Region of Paraguay. During this period, the Maká population residing in MR Alonso and Villa Hayes was estimated at 1700 people.(10, 11) Prisoners, health care workers (HCW), and patients with drug resistant TB were excluded because of the different risk profile for treatment outcome. Maká TB patients not living in MR Alonso or Villa Hayes were excluded due to regional differences in health services availability. Indigenous people from other communities were excluded as well. Furthermore, we excluded patients who died, were transferred or defaulted before the start of TB-treatment, whose treatment was still ongoing and patients with insufficient data. The study population was divided into the Maká and the non-indigenous population. (Figure 1.)

**Figure 1:**
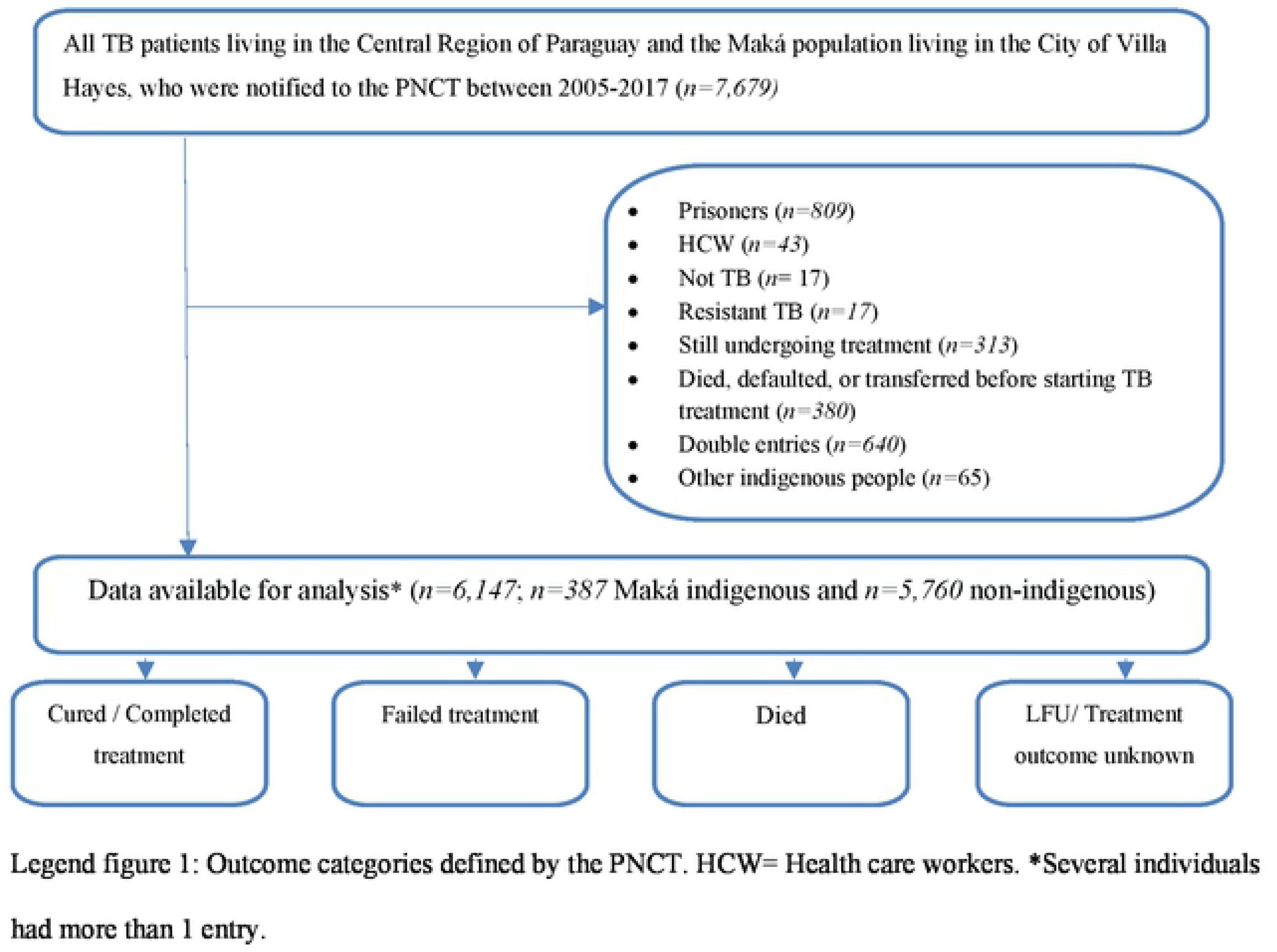
Inclusion and Exclusion criteria for study population.

### Data collection

Each Paraguayan region submits a monthly TB report to the PNCT with the number of new and recurrent cases and patient’s characteristics.(12) All reports since 2005 have been manually digitalized and are entered in an Excel database by their statistical department. To enable data analysis over the period 2005-2017, a new Excel file was made containing only the TB-cases that met the primary inclusion criteria. Double entries of recurrent cases were removed. Data quality of the digitalized dataset was assessed by cross checking the digital data with the original paper-based patient files for ~200 TB patients.

### Outcome definitions

TB treatment outcomes were categorized into ‘successful’ and ‘unsuccessful’ treatment outcome, applying the adapted WHO definitions used by the PNCT.

Successful treatment outcomes included the following categories:

- ‘cured’: positive status at start of treatment, completion of regimen, and at least two cases of negative smear microscopy, of which one at the end of treatment;
- ‘treatment completed’: positive status at start of treatment, completion of regimen with negative smear microscopy during control, but no final smear to ensure cure.

Unsuccessful TB-treatment outcome included the following categories:

- ‘treatment failure’: smear positive status after 5 months of treatment;
- ‘death’: death during TB treatment;
- ‘lost to follow-up’ (LFU): transfer out of the cohort during treatment or interruption of TB medication without clinical implications, for at least a month;
- ‘unknown’: patients without registered treatment outcome nor reason for LFU.

### Variables

Explanatory variables included socio-demographic and clinical characteristics. Age was categorized into 0-15, 16-32, 33-51, and >51 years of age. Place of diagnosis was defined as the ultimate source of the TB diagnosis report and included the following: the respiratory hospital INERAM in Asunción, specialized hospitals as defined by the ministry (indigenous hospital in Limpio), regional/district hospitals, private/social hospitals, medical dispensaries/general practices/health posts, or other/unspecified. TB was diagnosed either bacteriologically (smear microscopy, *Mycobacterium Tuberculosis* culture or GeneXpert), clinically, or unknown when no diagnostic method was registered. A person was defined as a ‘contact’ when he/she was a known contact of a TB patient. Admission categories were ‘new TB case’ or ‘previously treated’. Type of tuberculosis was either pulmonary (PTB), extra pulmonary (EPTB), or a combination of PTB/EPTB. Control smears were performed when one or more control sputum smear microscopy was registered during treatment. HIV-testing was defined as positive, negative, unknown, or not performed. Co-morbidities were categorized as substance abuse (alcohol, smoking or drug-use), Diabetes Mellitus, HIV/AIDS, multiple of these co-morbidities, other (cancer, asthma, renal failure, autoimmune disease, etc.), or unregistered.

### Statistical analysis

Tuberculosis incidence was calculated using the population numbers from the statistical department of the Paraguayan Ministry of Health. (11, 13) The indigenous patients from other communities were included in the total population/patient numbers, prisoners were excluded.

Demographics were analyzed separately for the Maká and the non-indigenous, and differences in the variables were assessed with the Pearson’s Chi-square test for categorical and Mann-Whitney U test for continuous variables. If a variable had more than 2 categories, the post-hoc Phi and Cramer’s V test was used to define the z-scores of each category. Age was analyzed both categorical and continuous (median and interquartile range [IQR]; no normal distribution).

To describe the performance of the TB registration system, a trend over time was assessed for several variables. Statistical significance was calculated using the Jonckheere-Terpstra test, comparing the proportion-distribution over the years.

All statistical analyses were done using IBM SPSS statistics (version 21.0; SPSS, Chicago, IL, USA), and GraphPad Prism (version 6.01, 2012; GraphPad Software, Cam USA). Statistical significance, unless stated otherwise, was assumed at *P*<0.05.

### Ethical considerations

The study was approved by the medical ethics committee of Paraguay (CEI-LCSP), with classification code 125/1104118, and carried out according to the principles of the Declaration of Helsinki and guidelines of the Council for International Organizations of Medical Sciences (CIOMS).(14)

## RESULTS

### TB incidence and patient demographics

The study population consisted of 6,147 TB patients, with 387 Maká (6.3%). (Figure 1.) The most important difference between the Maká and the non-indigenous population in the period from 2012 to 2016 was a 66 times higher incidence of the Maká (1,792 vs. 27 cases / 100,000 inhabitants). The Maká showed higher treatment success rates (75.2% vs. 67.8%; P=0.008), but a lower cure rate (8.7% vs. 36.6%; P<0.001). Both populations had around 22% of lost to follow-up (LFU).

Patient characteristics are described in Table 1. Of all TB-patients, 89.4% were new cases, and 10.6% (N=659) received previous treatment. The median age at diagnosis of the Maká was 3 years (IQR:1.3-19 years) and 51% were male, whereas the median age of the non-indigenous was 35 years (IQR:23-53 years), with 66% males. Maká were less often bacteriologically tested at diagnosis than the non-indigenous (55.0% vs. 77.8%; P<0.001), had higher DOTS coverage (89.4% vs. 47.1%; P<0.001), less control sputum smears (19.4% vs. 59.7%, P<0.001), less HIV testing (27.9% vs. 47.4%; P<0.001), and less registered co-morbidities (0.8% vs. 22.1%; P<0.001).

**Table 1:**
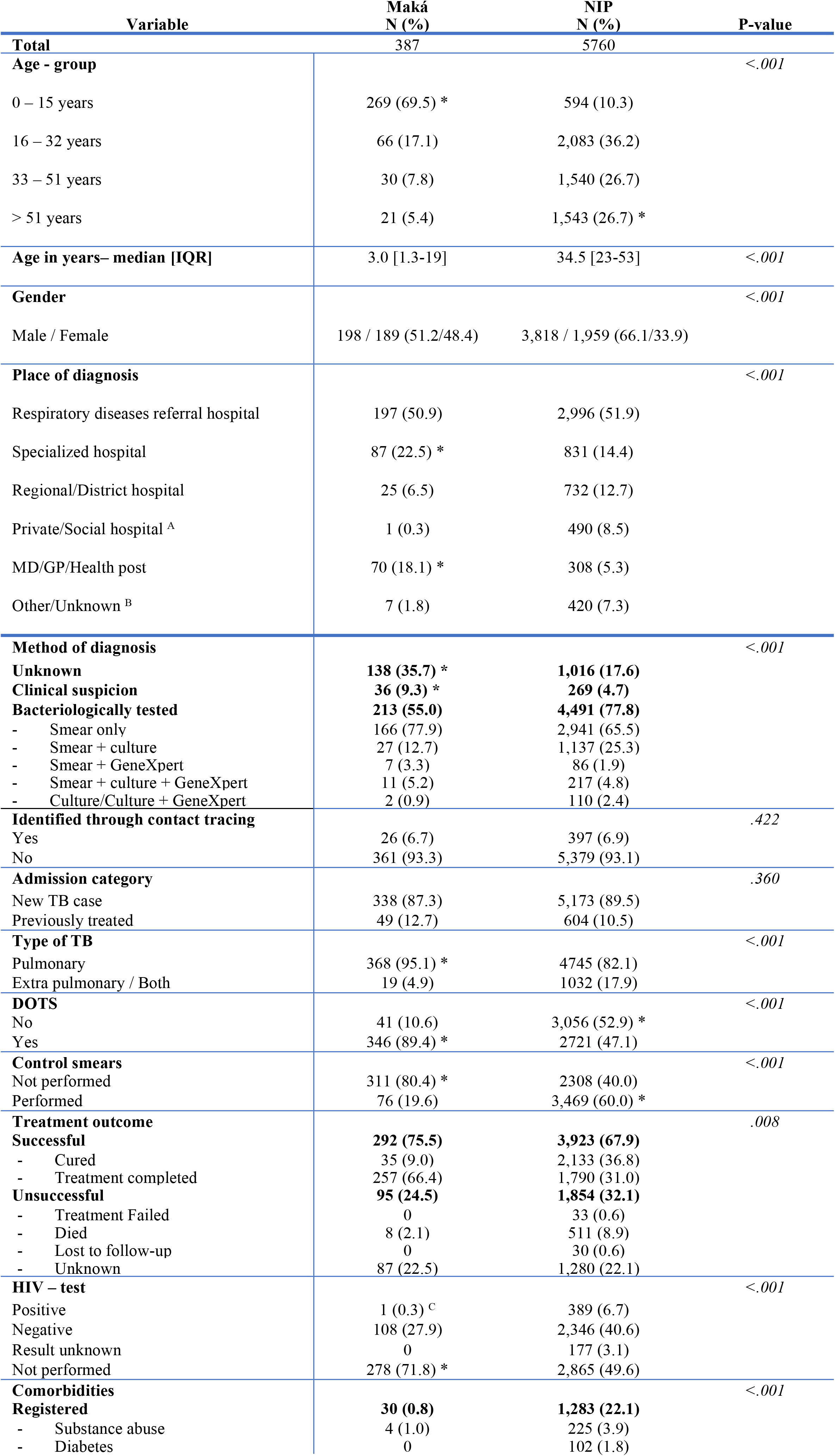
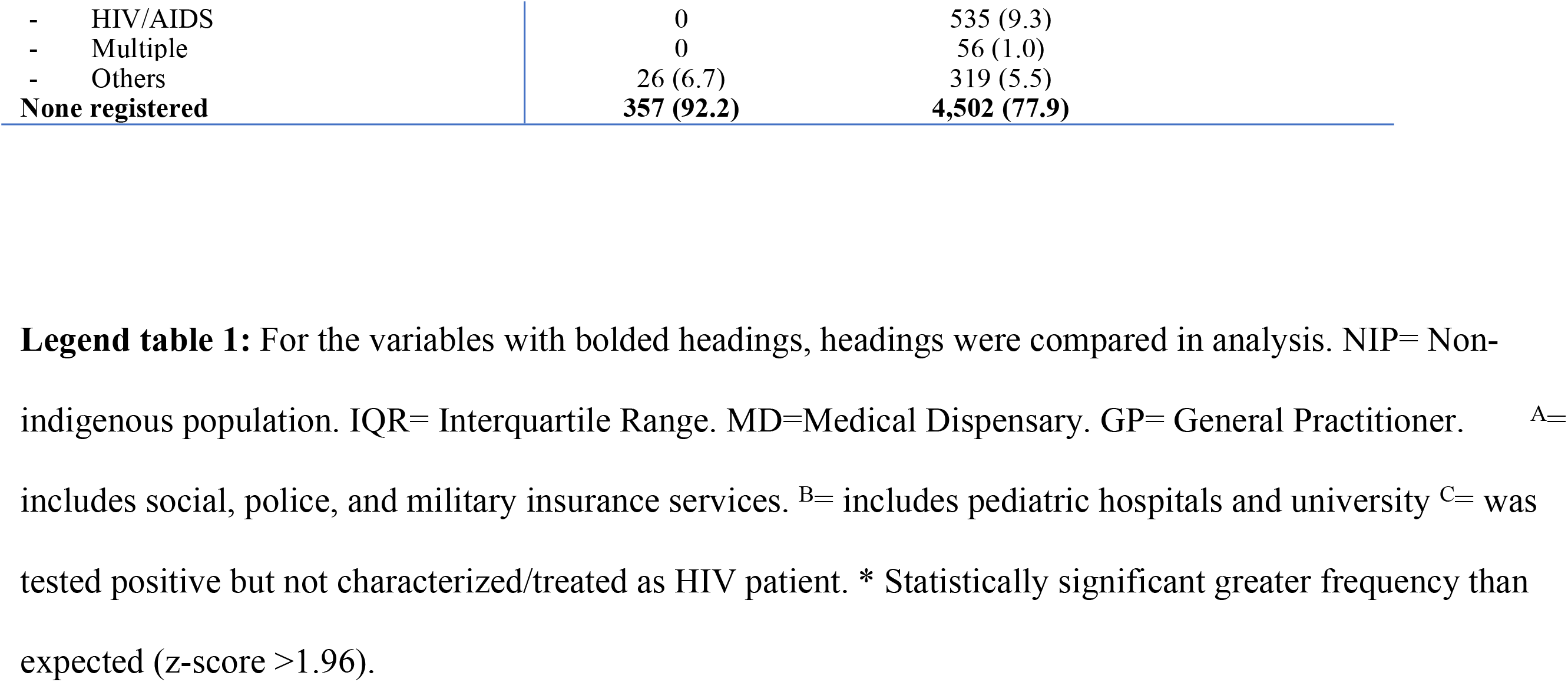
Study population characteristics of TB patients in the Central Region of Paraguay, diagnosed between 2005-2017.

| Variable | Maká<br>N (%) | NIP<br>N (%) | P-value |
| --- | --- | --- | --- |
| **Total** | 387 | 5760 |  |
| **Age - group** |  |  | <.001 |
| 0 – 15 years | 269 (69.5) * | 594 (10.3) |  |
| 16 – 32 years | 66 (17.1) | 2,083 (36.2) |  |
| 33 – 51 years | 30 (7.8) | 1,540 (26.7) |  |
| > 51 years | 21 (5.4) | 1,543 (26.7) * |  |
| **Age in years– median [IQR]** | 3.0 [1.3-19] | 34.5 [23-53] | <.001 |
| **Gender** |  |  | <.001 |
| Male / Female | 198 / 189 (51.2/48.4) | 3,818 / 1,959 (66.1/33.9) |  |
| **Place of diagnosis** |  |  | <.001 |
| Respiratory diseases referral hospital | 197 (50.9) | 2,996 (51.9) |  |
| Specialized hospital | 87 (22.5) * | 831 (14.4) |  |
| Regional/District hospital | 25 (6.5) | 732 (12.7) |  |
| Private/Social hospital <sup>A</sup> | 1 (0.3) | 490 (8.5) |  |
| MD/GP/Health post | 70 (18.1) * | 308 (5.3) |  |
| Other/Unknown <sup>B</sup> | 7 (1.8) | 420 (7.3) |  |
| **Method of diagnosis** |  |  | <.001 |
| **Unknown** | **138 (35.7) \*** | **1,016 (17.6)** |  |
| **Clinical suspicion** | **36 (9.3) \*** | **269 (4.7)** |  |
| **Bacteriologically tested** | **213 (55.0)** | **4,491 (77.8)** |  |
| - Smear only | 166 (77.9) | 2,941 (65.5) |  |
| - Smear + culture | 27 (12.7) | 1,137 (25.3) |  |
| - Smear + GeneXpert | 7 (3.3) | 86 (1.9) |  |
| - Smear + culture + GeneXpert | 11 (5.2) | 217 (4.8) |  |
| - Culture/Culture + GeneXpert | 2 (0.9) | 110 (2.4) |  |
| **Identified through contact tracing** |  |  | .422 |
| Yes | 26 (6.7) | 397 (6.9) |  |
| No | 361 (93.3) | 5,379 (93.1) |  |
| **Admission category** |  |  | .360 |
| New TB case | 338 (87.3) | 5,173 (89.5) |  |
| Previously treated | 49 (12.7) | 604 (10.5) |  |
| **Type of TB** |  |  | <.001 |
| Pulmonary | 368 (95.1) * | 4745 (82.1) |  |
| Extra pulmonary / Both | 19 (4.9) | 1032 (17.9) |  |
| **DOTS** |  |  | <.001 |
| No | 41 (10.6) | 3,056 (52.9) * |  |
| Yes | 346 (89.4) * | 2721 (47.1) |  |
| **Control smears** |  |  | <.001 |
| Not performed | 311 (80.4) * | 2308 (40.0) |  |
| Performed | 76 (19.6) | 3,469 (60.0) * |  |
| **Treatment outcome** |  |  | .008 |
| **Successful** | **292 (75.5)** | **3,923 (67.9)** |  |
| - Cured | 35 (9.0) | 2,133 (36.8) |  |
| - Treatment completed | 257 (66.4) | 1,790 (31.0) |  |
| **Unsuccessful** | **95 (24.5)** | **1,854 (32.1)** |  |
| - Treatment Failed | 0 | 33 (0.6) |  |
| - Died | 8 (2.1) | 511 (8.9) |  |
| - Lost to follow-up | 0 | 30 (0.6) |  |
| - Unknown | 87 (22.5) | 1,280 (22.1) |  |
| **HIV – test** |  |  | <.001 |
| Positive | 1 (0.3) <sup>c</sup> | 389 (6.7) |  |
| Negative | 108 (27.9) | 2,346 (40.6) |  |
| Result unknown | 0 | 177 (3.1) |  |
| Not performed | 278 (71.8) * | 2,865 (49.6) |  |
| **Comorbidities** |  |  | <.001 |
| **Registered** | **30 (0.8)** | **1,283 (22.1)** |  |
| - Substance abuse | 4 (1.0) | 225 (3.9) |  |
| - Diabetes | 0 | 102 (1.8) |  |

|  |  |  |
| --- | --- | --- |
| - HIV/AIDS | 0 | 535 (9.3) |
| - Multiple | 0 | 56 (1.0) |
| - Others | 26 (6.7) | 319 (5.5) |
| **None registered** | **357 (92.2)** | **4,502 (77.9)** |

**Legend table 1:** For the variables with bolded headings, headings were compared in analysis. NIP= Non-indigenous population. IQR= Interquartile Range. MD=Medical Dispensary. GP= General Practitioner. <sup>A=</sup> includes social, police, and military insurance services. <sup>B=</sup> includes pediatric hospitals and university <sup>C=</sup> was tested positive but not characterized/treated as HIV patient. \* Statistically significant greater frequency than expected (z-score >1.96).

Of the bacteriologically tested patients (4,707/6,147 = 76.5%), the Maká mostly had negative smears (62.4%), of which only a small percentage was further analyzed with culture and/or GeneXpert. 29 patients (13.6%) were culture and/or GeneXpert confirmed TB infections. The non-indigenous had 841 (18.7%) negative smears, of which 35.3% had further analysis with culture/GeneXpert. 1097 patients (24.4%) were culture/GeneXpert confirmed TB infections.

### TB patient-registration performance

The performance of TB patient-registration over the years is shown in Figure 2. In the period 2005-2016, there was a statistically significant trend of more HIV-tests, more sputum smear controls, less LFU cases, more smear microscopy, culture, and clinical diagnostic registration, and less patients with an unknown diagnostic method. From 2011-2016, the number of HIV tests, culture, GeneXpert and clinical diagnostic registration showed a statistically significant increase, while DOTS coverage and the number of unknown diagnostic methods decreased statistically significant.

**Figure 2:**
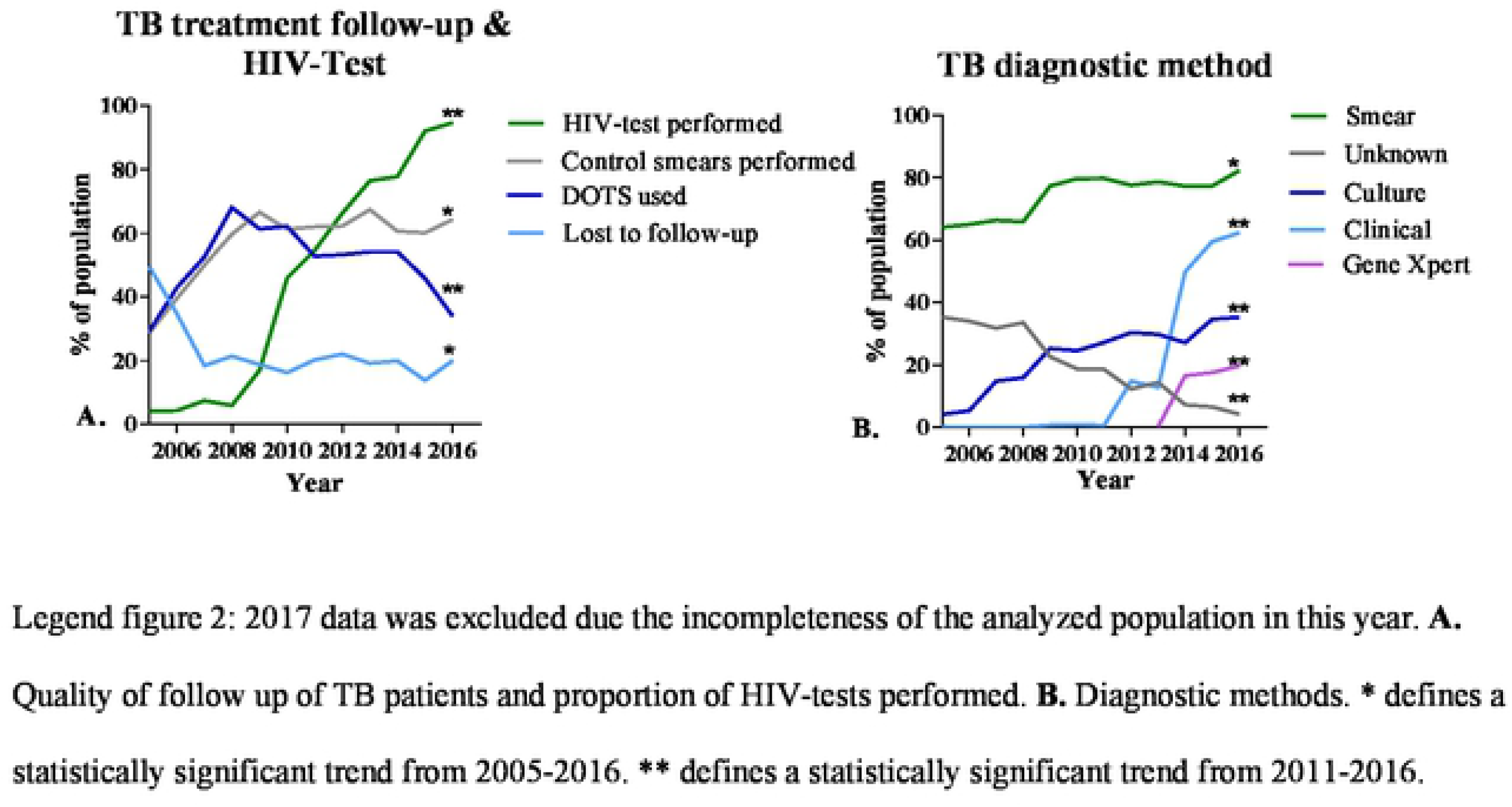
Performance of data collection of TB patients in the Central Region of Paraguay and the Maká population in Villa Hayes, analyzed by year of starting treatment and population proportion.

Of the LFU patients (including the ‘unknown’ outcome), 2.1% had a registered reason of which the most prevalent was (nomad) traveling. Treatment duration of LFU patients (both treated under DOTS and not) was not registered in 78%. In the ‘previously treated’ patient group, the Maká more often did not have information registered about their previous treatment, but this was not statistically significant (44.9% vs. 28.3%, P=0.257). The Maká had previous treatment failure in 4.1% and 10.2% abandoned treatment. None had given a reason for abandoning treatment. Of the non-indigenous population, 6.3% had had previous treatment failure, and 24.3% had abandoned treatment, of which 86 patients had decided to stop because ‘they felt recovered’. The percentage that had finished their previous treatment was similar.

The recurrence rates by TB treatment outcome are shown in Table 3. The recurrence rate was much higher for TB-patients with an unknown treatment outcome compared to a favorable treatment outcome and having an unknown treatment outcome increased the risk of a recurrence significantly (RR=23.7% vs. 5.7%. OR 5.15; 95% CI 4.32-6.14).

**Table 3:** Recurrence rate of TB patients living in the Central Region of Paraguay (N=6147).

| First treatment outcome: | Total | Patients with recurrent TB | Recurrence rate (%) | Recurrence risk (RR, 95% CI) | P-value |
| --- | --- | --- | --- | --- | --- |
| Favorable | 4,089 | 233 | 5.70 | *Ref* |  |
| Unfavorable | 523 | 51 | 9.75 | 1.79 (1.30-2.46) | <.001 |
| Unknown | 1,551 | 368 | 23.73 | 5.15 (4.32-6.14) | <.001 |

Legend table 3: Recurrence risk: difference in recurrence rate when having an unfavorable/unknown TB treatment outcome in the first treatment, compared to having a favorable first outcome.

## DISCUSSION

This study is the first to describe the incidence and treatment outcomes of the Maká in comparison to the non-indigenous population in Central Paraguay. While the Maká represent only ~0.1% of the total population in the Central Region of Paraguay, they comprised 6% of all TB reports in the Central Region within the period of analysis.

The TB incidence among the Maká (1,792/100,000 inhabitants) was 7 times higher than the incidence previously estimated by the National TB Program (272/100,000 inhabitants). Compared with the non-indigenous population the TB incidence among the Maká was even 66 times higher. This finding supports a study performed in Paraguay in 2003, which described a very high susceptibility to TB in the Ache indigenous population (3,700/100,000 inhabitants)(3), and a global systematic review in 2016 stated that the incidence of TB is generally higher in indigenous people.(15)

Overall, treatment success rates in the Central Region of Paraguay were below the WHO target of 85%.(16) The Maká had higher success rates than the non-indigenous population. The number of LFU patients was high for both populations and did not show significant improvement over the years. This implies the presence of an information gap, and leaves room for improvement of the patient registration and follow-up.

As the median age of the Maká was very young (3 years) and establishing TB diagnosis in children with sputum samples is challenging, the cure rates according to WHO definitions were difficult to obtain and therefore low.(17-19) The reason for higher treatment success rates of the Maká is probably explained by the higher ‘DOTS’ coverage of mothers taking care of their children.

The high prevalence of childhood TB in Maká raises two questions. Firstly; are these diagnoses in children robust? As TB diagnosis at young age is complicated it is possible that a part of these Maká are false positive TB diagnoses. In indigenous populations in Brazil, the same phenomenon in children has been described: one third of TB treatments were initiated without carrying out all diagnostic possibilities.(20, 21) In our study, even though the Maká children had more bacteriological testing compared to the non-indigenous, only a very small percentage had a confirmation with either a positive smear microscopy, culture or GeneXpert: 5.2% vs. 35.95% in the non-indigenous. (Figure 2.) Stigma in the Maká children could have led to overdiagnosis, as indigenous are known to be of higher risk for TB infection. This, together with the lack of clinical diagnosis, radiography and contact tracing indicates suboptimal and less reliable TB diagnoses in this group of patients which might have resulted in unnecessary treatments. Nevertheless, even with a substantial proportion of potentially false positive TB diagnoses, the TB incidence would probably still be higher among Maká compared to the non-indigenous population.

**Figure 3:**
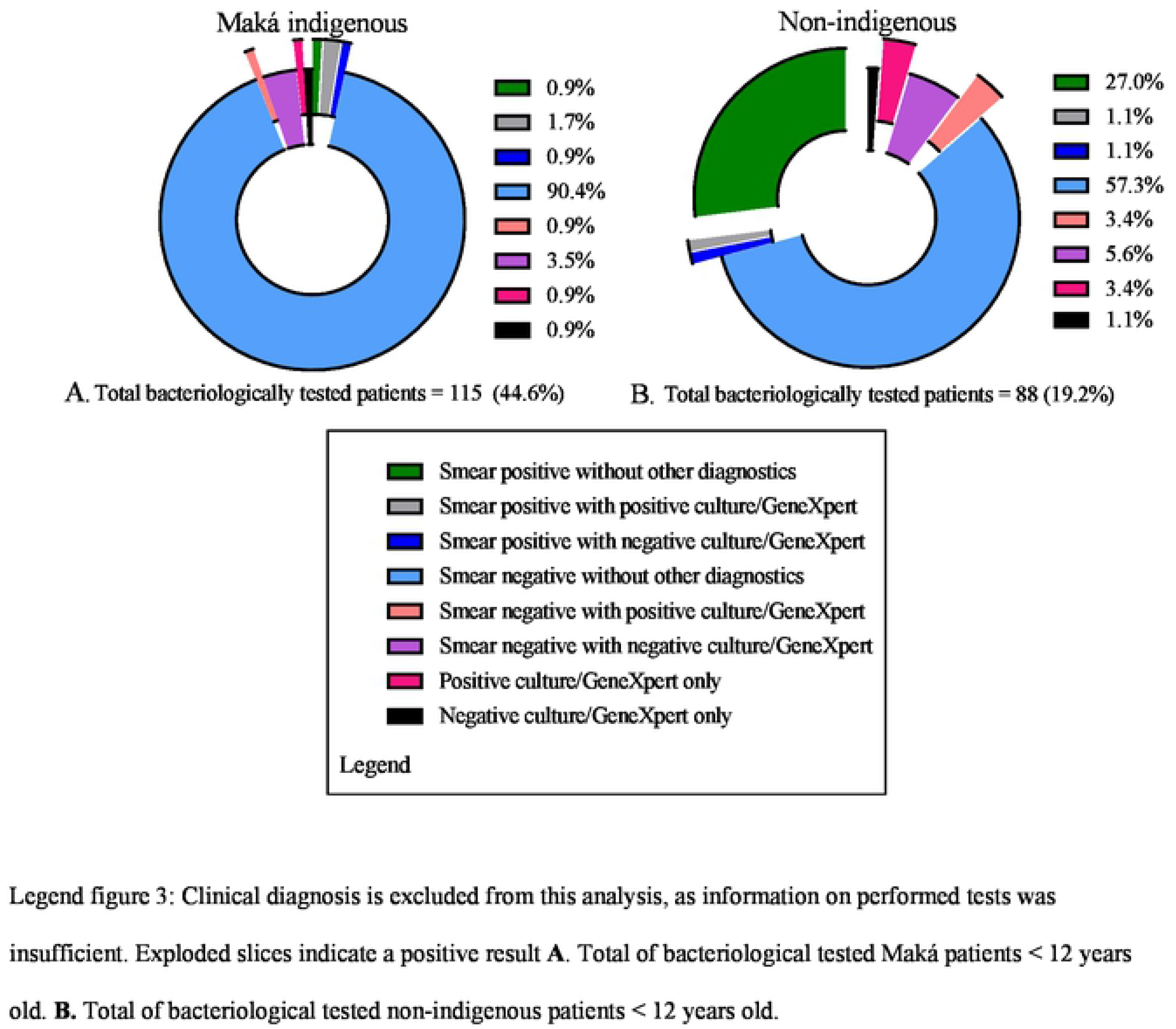
Performed diagnostics of the bacteriologically tested patients < 12 years old, in percentage of total.

Secondly, assuming a robust diagnosis of childhood TB cases; why is the incidence for Maká adults so low? Possibly there is a large reservoir of undetected adult pulmonary TB patients. Pediatric TB cases are generally considered less infectious and are most often caused by household contacts.(22-26) The Maká families live in single room houses (11, 27) with an increased risk of transmission, as sleeping with adults, crowded houses, and bad ventilation are known risk factors for pediatric infections.(22, 28-30) Additionally, the large number of unknown treatment outcomes implies that follow-up of TB patients is insufficient, which could lead to recurrence of disease and ongoing transmission.(7, 31)

This research has some limitations. We analyzed a manually digitalized dataset which may contain registration errors. The ~200 paper cross-checked files did not reveal major discrepancies, and therefore we do not expect that this manual digitalization would affect the study results. Nonetheless, errors might have occurred in the classification and/or diagnoses of the TB cases reported to the PNCT. Overdiagnosis of TB may cause an overestimation of the true TB burden in this population.

Furthermore, there were no annual numbers of the total population of the Maká and prisoners of the Central Region. Consequently, to calculate the TB incidence we worked with the population numbers that were present. To subtract the prisoners from the Central Region’s population, the *Censo Nacional 2013* was used, assuming that the number of incarcerated people remained stable over the years.(13) For the Maká, the general population growth of the Central Region was applied, using the Maká population number of the *Censo Nacional 2012* as a reference. This could mean that the TB incidence of the Maká is a slight overestimation, as their reproduction level is higher than the non-indigenous population’s.(11)

## CONCLUSION

This study provides a unique, detailed epidemiological description of TB cases in the Paraguayan Central Region. It showed that the incidence of pediatric TB in the Maká is extremely high, and the overall treatment success rate is below the WHO target of 85%. Furthermore, the study identified important differences between the indigenous and the non-indigenous population, regarding age, diagnostic methods, HIV-testing and DOTS coverage. Further exploration of the actual (childhood) TB burden in the Maká is necessary to obtain a trustworthy image of the population and guide TB-control measures. Assessment of the clinical diagnostics performed by checking the in-hospital registration forms might increase the likelihood and the reliability of the TB diagnoses among the Maká children. Mass screening of the Maká community could be an effective research method to determine the TB burden in the population, as well as identifying the potential “hidden” infectious reservoir that may be causing ongoing transmission. Bacteriological confirmation in this research is fundamental to achieve reliable data. Improved registration of the diagnostic methods and the follow-up data of the patients will enable the evaluation of the impact of TB-control interventions and more thorough research in the future.

## ACKNOWLEDGEMENTS

Many thanks to Natalia Sosa and Eva Chamorro of the National TB program for the help with the dataset and the background information on the National TB Program and the indigenous Maká population.

There was no conflict of interest.

## AUTHOR CONTRIBUTIONS

Conceptualization CM/AT/JF/GS/SA. Data Curation JF/GS/AT. Formal Analysis JF/GS. Funding Acquisition Not applicable. Investigation JF/ SA/CM. Methodology JF/GS/AT. Project Administration JF/CM. Resources SA/CM. Supervision CM/AT/GS. Validation JF/AT/GS. Visualization JF. Writing – Original Draft Preparation JF/CM. Writing – Review & Editing JF/AT/GS/SA/CM

